# Environmental DNA and wildlife camera traps uncover complimentary vertebrate visitation patterns at freshwater granite rock-holes

**DOI:** 10.1101/2024.10.31.621266

**Authors:** Brock Adam Hedges, Perry G Beasley-Hall, Tina E Berry, Kathryn L Dawkins, Andrew D Austin, Philip Weinstein, Michelle T Guzik

## Abstract

Freshwater ecosystems are in decline globally. In Australia, threatening processes include invasive species, increasing drought frequency, climate change and changes to land use, all of which have been associated with declining vertebrate diversity, particularly in Australia’s arid interior. Efficient monitoring tools are required to effectively monitor and conserve freshwater ecosystems and their associated vertebrate communities. Environmental DNA (eDNA) metabarcoding is one tool that shows promise for monitoring these systems, but knowledge of how eDNA data compares to more established ecological assessment techniques is limited. To address this knowledge gap, we sampled vertebrate eDNA from seven freshwater water bodies of proposed conservation importance in the Australian arid-lands, at three timepoints to measure visitation and compare our findings to camera trapping data at the same locations. Using eDNA we detected 19 species of vertebrates, including native species (such as macropods, wombats and emus) and invasive species (such as feral goats, cats and foxes). In contrast, camera traps detected 32 species, and was much more successful at detecting bird visitation than eDNA. These communities varied both spatially between rock-holes, and temporally, with summer collection periods being distinct from winter-spring. Our results demonstrate the success of eDNA metabarcoding as a tool for monitoring vertebrate visitation to arid-lands freshwater ecosystems that is complementary to more traditional survey methods such as wildlife camera trapping. Finally, we provide conservation recommendations for these vertebrate communities and discuss the efficacy of eDNA for monitoring freshwater resources in arid-lands environments.

## Introduction

Freshwater ecosystems and their riparian zones are experiencing some of the most rapid habitat declines globally and the rate of decline has increased in recent decades (Albert et al. 2021). Most impacts can be attributed to human activities including changes in land use, pollution, and the effect of climate change (Albert et al. 2021, Mulero et al. 2021). Freshwater ecosystems are of especially high value in semi-arid and arid regions due to the relative scarcity of accessible surface water (Noy-Meir 1973, Porporato et al. 2002), and correspondingly are amongst the most vulnerable ecosystems in the face of environmental change. In such regions, small bodies of water can occur but are typically ephemeral, being only sporadically wet throughout the year due to their reliance on rainfall (Bayly 1997, Bayly 1999b, Bayly 2001). For declining freshwater habitats, potential recovery in some cases and slowing of losses in others is possible but these are likely to require coordination of efforts at local, regional, national, and international levels (Albert et al. 2021, IPCC 2022b). For such measures to be devised and enacted, it is essential that biodiversity reliant on these habitats is surveyed and monitored for improved decision making and management.

In Australia, droughts are increasing in both frequency and severity due to climate change (Ndehedehe et al. 2021) and are projected to continue to do so under all future emissions scenarios (IPCC 2022a). Freshwater ecosystems are particularly vulnerable due to their dependence on thermal and hydrological regimes and climate change impacts (Filipe et al. 2012, da Silva et al. 2023). For arid-lands ecosystems where fauna are dependent on scarce freshwater resources, targeted monitoring programs with long-term data are rare in Australia and accurate distribution and abundance data is frequently lacking or non-existent for many declining taxa (Bino et al. 2020, Scheele et al. 2019).

Monitoring effort is also hampered by accessibility to remote areas, as well as their sparsely populated nature and a lack of funding. For these reasons, landscape-wide monitoring tools are required that will allow for cost-effective determination of remote freshwater ecosystems, biotic community health, and characterisation of the resource value to Australian terrestrial fauna (Beasley-Hall et al. 2023).

The security of Australia’s unique vertebrate biodiversity is of growing concern (Bradshaw 2012, Bilney 2014, Hanna and Cardillo 2014, Woinarski et al. 2015, Geyle et al. 2018). Australia has experienced a native mammal extinction crisis since European colonisation, with terrestrial mammal losses accounting for >10% of species lost over the last 200 years (Woinarski et al. 2015). A third of all mammal extinctions after the year 1500 CE have occurred in Australia (Fisher et al. 2014). The loss of small to medium-sized species in Australia especially has led to a decline in particular taxonomic groups, many of which are endemic to arid regions. In particular, mammals within the ‘critical weight range’ of 0.035 – 5.5 kg account for the vast majority of these extinctions (Short and Smith 1994, Johnson and Isaac 2009, Moseby et al. 2009, Murphy and Davies 2014). Mammalian losses have also impacted various ecosystem services, such as bioturbation by small burrowing mammals (Fleming et al. 2014) and predation by marsupials (Moseby et al. 2021). Australia’s bird fauna is also considered to be in decline, with multiple extinctions having occurred since European colonisation (Berryman et al. 2024, Woinarski et al. 2024), and more likely to occur over the next two decades (Campbell et al. 2024, Geyle et al. 2018). Whilst the extent to which Australia’s reptile fauna is similarly imperilled is not as well-known due to data deficiency (Geyle et al. 2021, Tingley et al. 2019), there is growing concern that extinction events will increase in frequency over the coming century (Geyle et al. 2021, Tingley et al. 2019).

Invasive predators (such as feral cats and foxes) are often considered to be the primary drivers of Australian small mammal decline (Burbidge and Manly 2002, Kutt 2012, Fisher et al. 2014, Frank et al. 2014, Hanna and Cardillo 2014), although environmental change (McKenzie et al. 2007), non-predatory invasive species (Woinarski et al. 2011), and epizootic diseases (Abbott 2006) are also contributors. Similarly, predation by invasive species, changes to habitat availability and complexity, and climate change are thought to be contributing to both bird and reptile decline (Campbell et al. 2024, Geyle et al. 2021, Senior et al. 2021, Woinarski et al. 2024). Whilst the global conservation effort to preserve vertebrate biodiversity is currently insufficient (Butchart et al. 2010, Hoffmann et al. 2010), some success has been achieved in Australia. For example, fenced exclosures (fenced areas designed to keep invasive or overabundant species out) in arid regions have improved survival of numerous threatened and reintroduced species in small pockets across the landscape (Moseby et al. 2009). A clear need exists to understand the changes in mammalian community composition, both native and invasive in arid-lands to inform management of populations, especially in the face of substantive impacts from climatic change. To date, monitoring efforts in these regions are frequently limited in their scope, and often target specific taxa rather than communities at a broader scale (Moseby et al. 2009, Beasley-Hall et al. 2023).

Monitoring methods for vertebrates can be biased and inconsistent, in part due to problems associated with transferability and repeatability, and standardisation of sampling techniques between projects and habitat types (Cross et al. 2020, McDonald et al. 2021). There is a clear need for interchangeable tools that allow for comparison between disparate programs (Beasley-Hall et al. 2023, Zhang et al. 2023). Environmental DNA (eDNA) metabarcoding has great potential as a transferable biodiversity monitoring tool across a range of ecosystems. Metabarcoding approaches represent a high-throughput DNA sequencing (HTS) technique that allows for the identification of multiple taxa from a single sample (Miya et al. 2020). This approach can characterise entire communities in addition to targeting specific species. Environmental DNA can be used as a standardised and repeatable approach which shows promise as a method for landscape or national level biomonitoring efforts (Sales et al. 2020a, Miya et al. 2020). Environmental DNA metabarcoding is particularly useful for detecting biodiversity from freshwater substrates irrespective of location and species present (e.g., Kuehne et al. 2020, White et al. 2020, West et al. 2021), as well as identifying abiotic stressors (Fan et al. 2020). Inference from these data can facilitate informed conservation management, including early detection of invasive animal and plant species to inform targeted removal and/or control (Kuehne et al. 2020).

To date, freshwater eDNA metabarcoding studies have focussed predominantly on species such as fishes (Hänfling et al. 2016), amphibians (Valentini et al. 2016), and macroinvertebrates (Johnsen et al. 2020, Klymus et al. 2020, Rodgers et al. 2020). However, these methods also show promise for the detection of terrestrial vertebrates that have recently interacted with relevant water bodies (McDonald et al. 2023). Successes in the detection of aquatic, semi-aquatic, terrestrial reptiles (West et al. 2021), and terrestrial mammals (Harper et al. 2019, Sales et al. 2020b) indicate freshwater eDNA metabarcoding is viable and comparable to traditional vertebrate survey methods (i.e., wildlife camera trapping, field surveys, etc.) per unit effort, with the potential to improve information on distribution of mammals at a landscape level (Sales et al. 2020b). For example, eDNA biomonitoring at inland springs has detected more terrestrial vertebrates than were observed with wildlife cameras, in person observations, or from indirect evidence (e.g., scats or tracks) (Parker et al. 2021). Whilst the presence of a species detected by eDNA is difficult to link to specific behaviours or activities (Harper et al. 2019), it is likely that visitation frequency should drive eDNA signal (Ushio et al. 2017), particularly in regions with low rainfall and for water bodies with low connectivity, as is the case for many freshwater habitats in the Australian arid-lands.

Freshwater eDNA metabarcoding performs well compared to traditional monitoring and detection techniques (Piggott et al. 2021). However, some well-established limitations exist for eDNA biomonitoring. For example, in natural ponds, terrestrial vertebrate eDNA is known to be distributed more unevenly than in experimental artificial water bodies (Harper et al. 2019), which may limit confidence in results where replication is limited. Further, eDNA biomonitoring is currently inadequate for estimating abundance and biomass (Di Muri et al. 2020). Finally, false negatives are also problematic (Sales et al. 2020a, Sales et al. 2020b). Whilst these limitations are fundamentally problematic and acknowledged in the literature as areas requiring further research, eDNA metabarcoding is still an emerging field that is overwhelmingly useful as a complementary method in assessing difficult to survey populations (Mas-Carrió et al. 2022). As such, freshwater eDNA metabarcoding is likely better used alongside more traditional surveying methods (Sales et al. 2020b); a combined approach that has only recently been considered (Farrell et al. 2022, Johnson et al. 2023).

Based on previous research that has demonstrated eDNA metabarcoding can be used to 1) detect vertebrate communities characterised as visitors to freshwater systems (Harper et al. 2019, Lyet et al. 2021), 2) can corroborate findings from more traditional methods of biomonitoring, such as camera trapping (Harper et al. 2019, Leempoel et al. 2020, Lyet et al. 2021, Mas-Carrió et al. 2022, Sales et al. 2020b) and 3) has proven effective in detecting variation in visitation rates with a variety of sampling methodologies (Mcdonald et al. 2023), here, we aimed to compare the efficacy of biomonitoring vertebrate fauna in the well-known but understudied freshwater habitats in arid Australia between eDNA metabarcoding and camera trapping methods. We specifically focused on the detection of vertebrates in the iconic arid-lands granite rock-holes at Hiltaba Nature Reserve. In doing so, we investigated the applicability of eDNA metabarcoding as a conservation management tool. Using a targeted wildlife camera trapping dataset we explore the complementary application of eDNA metabarcoding and this traditional technique at Hiltaba Nature Reserve; a notable location of higher-than-average habitat quality in the Australian arid-lands.

Our specific aims were to a) test the application of eDNA metabarcoding in detecting native and invasive vertebrate visitation to the Australian arid-lands freshwater granite rock-holes system, b) compare detection of vertebrates with eDNA metabarcoding to conventional wildlife camera trapping techniques, c) determine whether vertebrate communities detected using eDNA varied spatially and temporally, and d) make a series of conservation and management recommendations for the freshwater granite rock-holes. We predicted an overlap between eDNA methods used here and traditional camera trapping methods with respect to the detection of native and invasive vertebrate communities. We also predicted that eDNA metabarcoding would be sensitive enough to detect spatial and temporal community variation.

## Methods

### Study site

The present study was conducted at Hiltaba Nature Reserve (HNR), a large ex-pastoral property that borders the Gawler Ranges National Park to the north of South Australia’s Eyre Peninsula (Figure 1). Approximately 78,000 hectares in area, HNR has been managed for conservation outcomes by the Nature Foundation (South Australia) since its acquisition by the organisation in 2012. Primarily composed of a series of rolling granite hills interspersed with woodland and grassland, HNR constitutes many relictual habitats utilised by local plants and animals known to be in decline elsewhere (Nature Foundation 2023). The Nature Foundation has enacted a series of conservation programs to improve biodiversity management at the reserve including destocking of pastoral species (primarily sheep), and the routine culling of invasive species of known ecological harm (primarily feral goats). We selected HNR due to its relative high quality as an ecological refuge compared to adjacent partly degraded habitat and consider it as a location optimal for ecological research.

**Figure 1.**
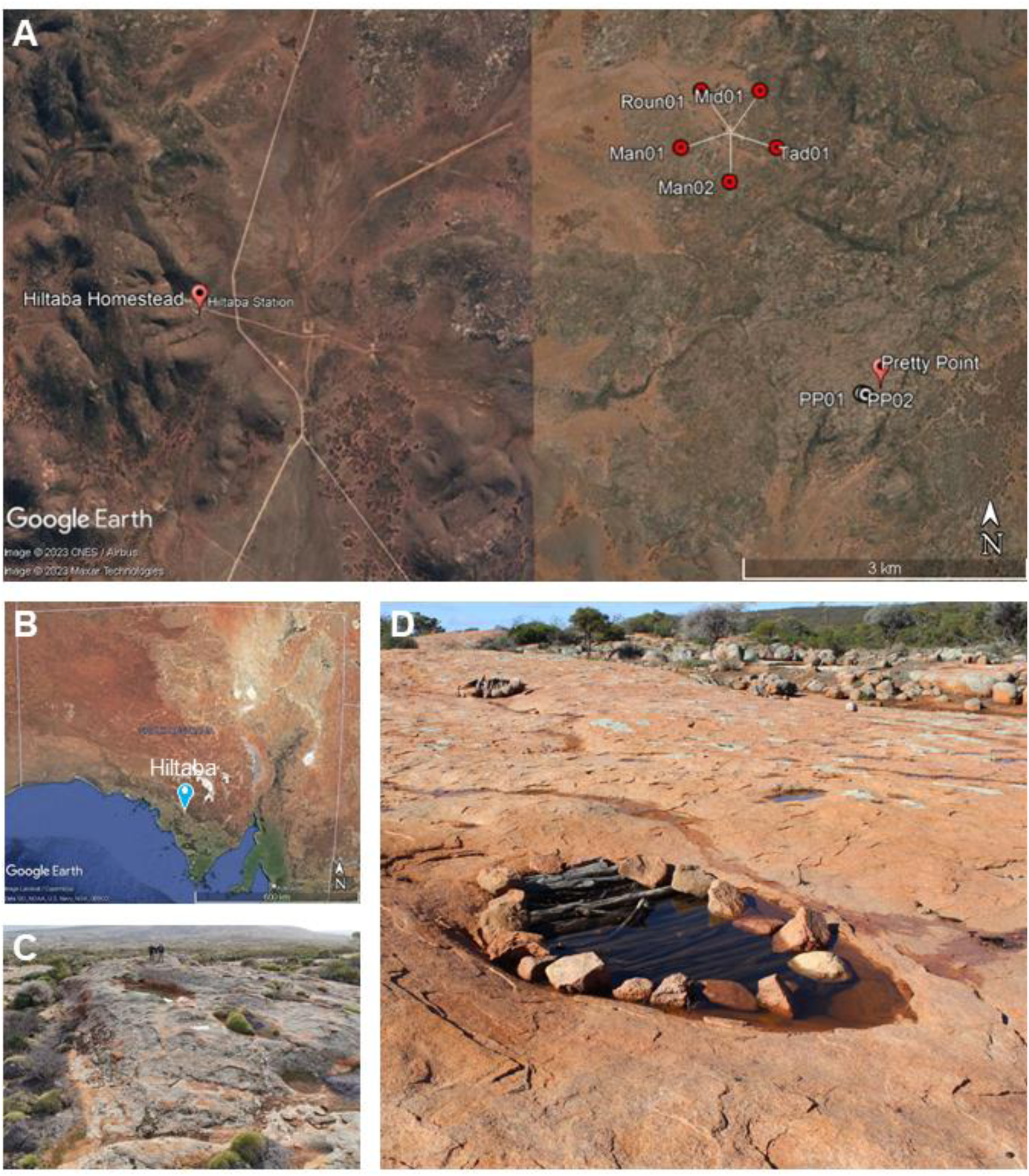
A) The locations of the seven rock-holes sampled for freshwater eDNA during 2020; B) the location of Hiltaba Nature Reserve (HNR) in South Australia; C) a rain-filled granite rock-hole at the ‘Pretty Point’ outcrop at HNR; D) a rain-filled granite rock-hole on the ‘photopoint outcrop’ at HNR that is actively managed by members of the Gawler Ranges Aboriginal Corporation, diameter of the rock-hole at widest point = 2.2 m.

The granite hills throughout much of the Reserve are often entirely exposed, forming granite outcrops which can serve as impermeable locations for storage of water and sediments at depressed points. These depressions or ‘rock-holes’ provide habitat for a suite of aquatic invertebrates and plants. As many vertebrate species have been observed visiting these rock-holes as a source of accessible freshwater, the granite rock-holes at HNR have been identified by the Nature Foundation as a location of critical conservation interest with respect to local vertebrate biodiversity. Notably, a subset of the rock-holes at HNR are managed using traditional cultural techniques by members of the Gawler Ranges Aboriginal Corporation, using practices that date back thousands of years prior to European occupation of Australia (Jenkin et al. 2011). These practices involve the use of timber and rocks to prevent animals from falling into the water, and regular cleaning (see Figure 1D).

### eDNA sampling

Water was sampled from five rock-holes in February 2020, and seven rock-holes in each of July and October 2020 (Table 1). Five 1 L replicates were collected from each rock-hole. One negative control was implemented for each rock-hole using 1 L of bottled water. Replicates were collected using 1 L wide mouth bottles (NALGENE^TM^). During February, only sites at outcrop 1 were sampled due to inaccessibility of rock-holes at outcrop 2 during the hot summer months. During October, rock-holes Man02 and PP02 were not sampled due to low water levels. Samples were collected whilst wearing disposable latex gloves, stored and transported in clean plastic tubs to minimise contamination.

**Table 1.** Granite rock-holes sampled for freshwater eDNA during February, July, and October trips (green checkmark). Rock-holes not sampled (red cross) were either empty or had only a small volume of water present. Rock-hole type assignments adapted from Timms (2013b).

| Rock-hole features |  |  |  |  |  | Sample period |  |  |
| --- | --- | --- | --- | --- | --- | --- | --- | --- |
| ID | Latitude | Longitude | Elevation (m) | Type | Outcrop | Feb | July | Oct |
| Man01 | -32.13941 | 135.13461 | 216 | Pit | 1 | ✓ | ✓ | ✓ |
| Man02 | -32.13943 | 135.13478 | 216 | Pit | 1 | ✓ | ✓ | ✗ |
| Tad01 | -32.1395 | 135.1348 | 216 | Pit | 1 | ✓ | ✓ | ✓ |
| Roun01 | -32.13931 | 135.13492 | 217 | Pan | 1 | ✓ | ✓ | ✓ |
| Mid01 | -32.13941 | 135.13461 | 217 | Pit | 1 | ✓ | ✓ | ✓ |
| PP01 | -32.165521 | 135.1488 | 312 | Pan | 2 | ✗ | ✓ | ✓ |
| PP02 | -32.165521 | 135.1488 | 310 | Pan | 2 | ✗ | ✓ | ✗ |

Sampling bottles were cleaned first with bleach solution, and then ethanol before reuse. Negative field controls (prepared in the field but using RO water) and equipment controls (prepared in the filtering room using RO water) were used to test for contamination.

Replicates and blanks were filtered through glass-fibre membranes with 0.44 µm pores using a vacuum pump (JAVAC, model CC-45) connected to a series of conical flasks. The first flask was filled with silica beads and the second was attached to a magnetic filter funnel. For the July and October samples, a Pall Sentino microbiology pump was used. A pore size of 0.44 µm was selected due to the high concentration of suspended solids present in rock-hole freshwater samples. Pump equipment was wiped down with 10% bleach solution and then ethanol between filtering for each rock-hole.

Membranes were stored on ice while in the field before being transferred to -20°C storage pending DNA extraction.

### eDNA laboratory methods

DNA was extracted from half of each filter paper using a modified Qiagen DNeasy Blood and Tissue Kit and an automated QIAcube extraction platform (QIAGEN, Germany). Where more than one filter paper was used for a sample, equal portions of each paper were taken to total a half filter paper. All extractions were undertaken in a dedicated PCR-free laboratory and extraction controls were processed alongside samples. Extractions were eluted in a final volume of 100 µL AE buffer (10 mM Tris-Cl, 0.5 mM EDTA).

To determine the required dilution for optimal amplification, PCR reactions were performed in duplicate on each extraction by adding DNA template directly to the neat PCR master mix then performing a serial dilution (1:10). The PCRs were performed at a final volume of 25 µL where each reaction comprised of: 1× PCR Gold Buffer (Applied Biosystems), 0.25 mM dNTP mix (Astral Scientific, Australia), 2 mM MgCl_2_ (Applied Biosystems), 1U AmpliTaq Gold DNA polymerase (Applied Biosystems), 0.4 mg/mL bovine serum albumin (Fisher Biotec), 0.4 µM proprietary forward and reverse primers, 0.6 μl of a 1:10,000 solution of SYBR Green dye (Life Technologies), and 2 µL template DNA. PCRs were performed on StepOne Plus instruments (Applied Biosystems) with the following cycling conditions: 95°C for 5 min, followed by 50 cycles of: 95°C for 30 sec, 49°C for 30 sec, 72°C for 45 sec, then a melt-curve analysis of: 95°C for 15 sec, 60°C for 1 min, 95°C for 15 sec, finishing with a final extension stage at 72°C for 10 min.

After selection of the optimal dilution (neat or 1:10), PCRs were repeated in duplicate as described above but instead using unique, single use combinations of 8 bp multiplex identifier-tagged (MID-tag) primers as described in Koziol et al. (2019) and van der Heyde et al. (2020). Master mixes were prepared using a QIAgility instrument (QIAGEN) in an ultra-clean lab facility, with negative and positive PCR controls included on every plate to ensure the validity of results. A sequencing library was created by combining samples into mini-pools based on the PCR amplification results from each sample. The mini-pools were then analysed using a QIAxcel (QIAGEN) and combined in roughly equimolar concentrations to form libraries. Libraries were then size selected (250 - 600 bp cut-off) using a Pippin Prep instrument (Sage Sciences) with 2% dye-free cassettes, cleaned using a QIAquick PCR purification kit, quantified on a Qubit (Thermo Fisher), and diluted to 2 nM. The libraries were sequenced on an Illumina MiSeq instrument using a 500-cycle V2 kit with custom sequencing primers.

### Bioinformatics and statistical analyses

Processing and analysis of eDNA sequence data was performed with QIIME2 v.2021.11 (Bolyen et al. 2019). Raw sequences were demultiplexed, manually trimmed by quality scores, filtered to remove chimeric sequences, and denoised using DADA2 (Callahan et al. 2016). Sequences were then tabulated to construct 100% OTUs and representative sequences were derived for each OTU. Representative sequences were aligned using MAFFT v7.511 (Katoh et al. 2002) and a midpoint-rooted phylogeny was generated using FastTree 2 for downstream calculation of diversity metrics (Price et al. 2010). A custom database was constructed using BLAST from vertebrate *16S* rRNA sequences available via the NCBI GenBank repository as of June 2022 using the following search query: (("16S ribosomal RNA" OR "16S rRNA" OR mitochondrion) AND Vertebrata[Organism]) NOT Homo sapiens[Organism]. BLASTN (Camacho et al. 2009) was then used to query our representative sequences against the custom database with the maximum number of target sequences set to 1 per query. Representative sequences with non-vertebrate top hits, or those without a top hit with a corresponding species-level assignment, were excluded from further analyses. Alpha diversity metrics were then calculated including evenness (Pielou 1966), Faith’s phylogenetic diversity (Faith 1992), and Shannon’s diversity index (Shannon and Weaver 1949). Beta diversity analyses were examined including Bray-Curtis (Bray and Curtis 1957), Jaccard (Jaccard 1901), and weighted and unweighted Unifrac distances (Lozupone et al. 2011) against metadata columns including rock-hole location, depth, and collection month. PCoA plots of these metrics were exported to R version 4.2.2 (RCoreTeam 2013) using *QIIME2R* v0.99.6 for further visualisation (Bisanz 2018). Final PCoA plots were generated using *ggplot2* v3.4.0 (Wickham 2016). Rarefaction curves were generated using *iNext* v3.0.0 (Hsieh et al. 2022).

### Wildlife camera trapping data

Motion-triggered wildlife cameras (Browning Dark Ops Pro XD) were deployed at each of the seven rock-holes, strapped to star-droppers in bare soil adjacent to each outcrop. Cameras were arranged approximately 1 m away from the outcrop edge, and 1 m above the ground. The cameras were deployed between 8–11 July 2020, and were collected between 8–9 June 2021. Wildlife cameras were programmed to take a high-resolution photograph immediately following movement sufficient to trigger an infra-red sensor. Visitation was recorded as a series of images from which vertebrate species could be identified. This methodology was reviewed and approved by the University of Adelaide animal ethics committee (see acknowledgements for details). Images were processed using the MegaDetector machine learning algorithm (Microsoft AI for Earth 2020) desktop application v0.0.2 (Gyurov 2022), and resulting outputs were identified manually to species (where possible).

## Results

### Environmental DNA

We successfully recovered vertebrate eDNA from 87% (79/91) of samples using the *16S* rRNA assay. Sequencing yielded 9,107,435 individual reads in total with an average of 90,173 sequences per sample. A total of 19 unique vertebrate taxa were detected, including 11 mammals, five birds, three reptiles, and two amphibians (Figure 2).

**Figure 2.**
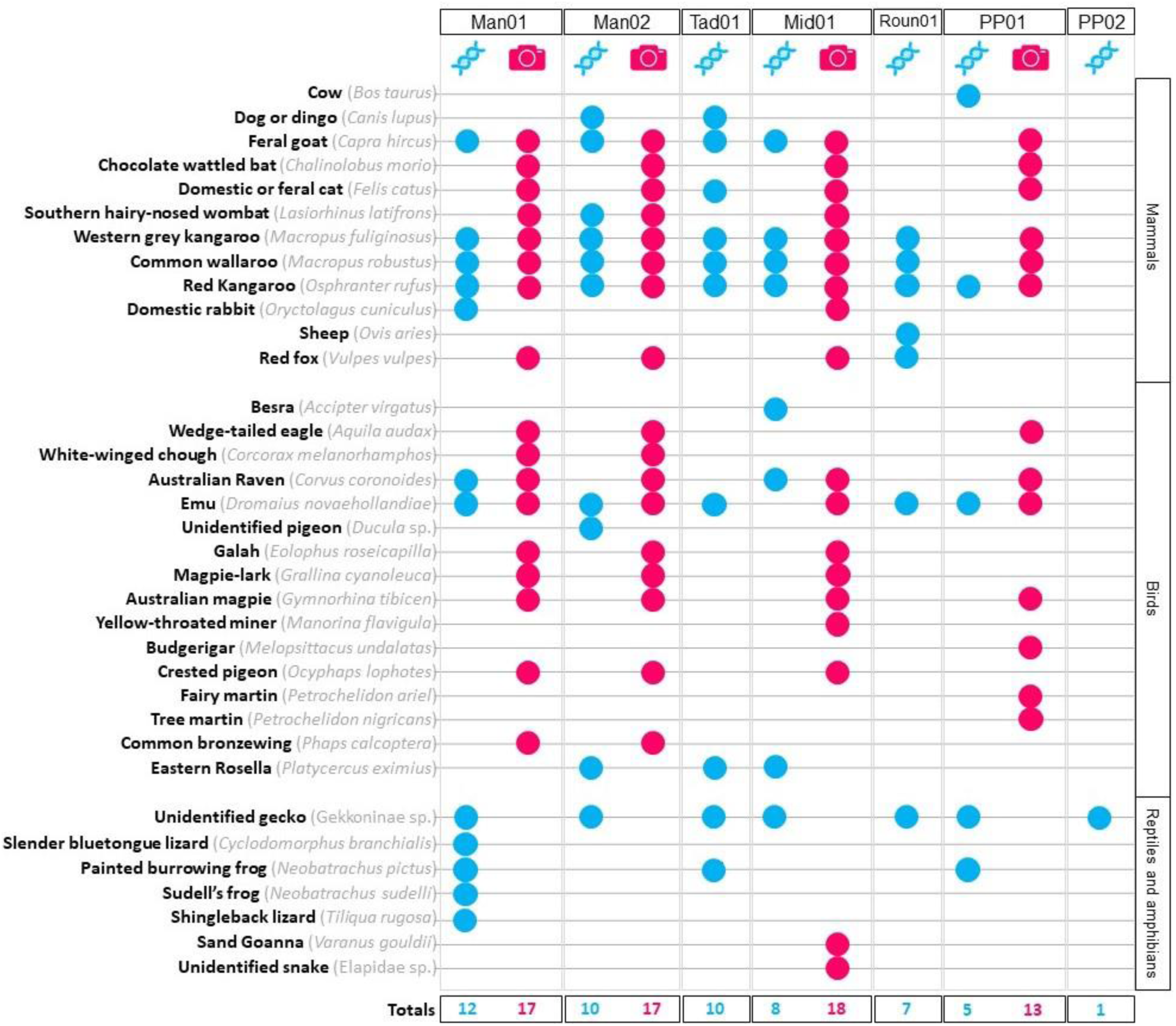
A comparison of the vertebrate taxa identified from freshwater eDNA collected from seven rock-holes at HNR throughout 2020 and with wildlife camera traps at the same rock-holes as presented in Hedges (2023). Blue circles indicate when a taxon was observed in the eDNA dataset, red circles indicate when a taxon was observed in the wildlife camera trap dataset. The complete dataset of presence absence records can be seen in supplementary table 1, and a comparison between eDNA data, camera trapping data from Hedges (2023) and bush blitz survey data from Commonwealth of Australia (2015a) can be seen in supplementary table 2.

Overall, eDNA metabarcoding was successful in recovering 54% of the species that had been detected using wildlife camera traps (19/35) (Figure 2). We recovered 91% of the mammal species we observed using wildlife cameras (11/12), with three new mammal records using eDNA that were not recorded using wildlife cameras (Figure 2). We also detected five reptile species that were not recorded with wildlife cameras (though they had been detected at rock-holes not surveyed with eDNA) (Hedges 2023) and generated two new reptile records and two new amphibian records not previously associated with rock-hole visitation. The application of eDNA was less effective for birds, with less than 11% of species recorded by camera traps in Hedges (2023) also having been recorded with freshwater eDNA, and three new records.

Rarefaction plots suggest that the number of rock-holes sampled was insufficient to capture 90% of the total vertebrate taxa that attend rock-holes at HNR (Figure 3A). However, within each rock-hole and at each timepoint rarefaction curves varied, with five replicates shown to be sufficient to capture >90% for some rock-holes and timepoints (Man01 in July, Man02 in Feb and July, Mid01 in Oct, PP01 in Feb and Round01 in Oct), whilst insufficient to capture 90% of taxa at others (Figure 3B).

**Figure 3.**
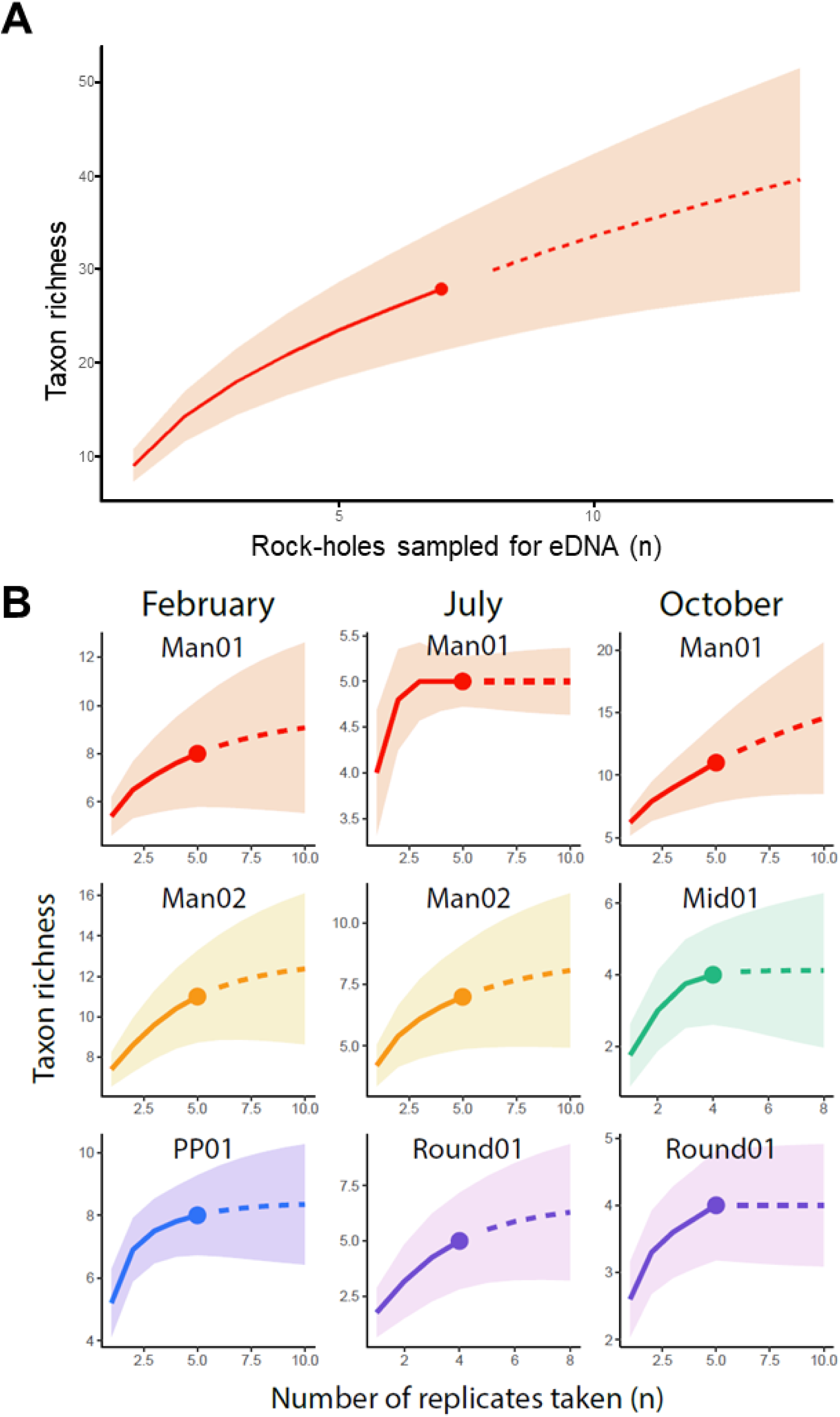
Species discovery curves showing A) accumulation of new records with each added rock-hole. Asymptote indicates that seven rock-holes was sufficient to recover 95 % of OTUs that are recoverable in filtered freshwater eDNA samples, and B) accumulation of new records with each added replicate. Asymptote indicates that five replicates plus one blank is sufficient to recover 95 % of OTUs that are recoverable in filtered freshwater eDNA samples

PCoA plots showing maximum dissimilarity for vertebrate occurrence between samples showed that samples collected in July and October were more similar to one another than either were to samples collected in February (Figure 4A). However, communities detected from individual rock-holes were generally distinct from one another (Figure 4B).

**Figure 4.**
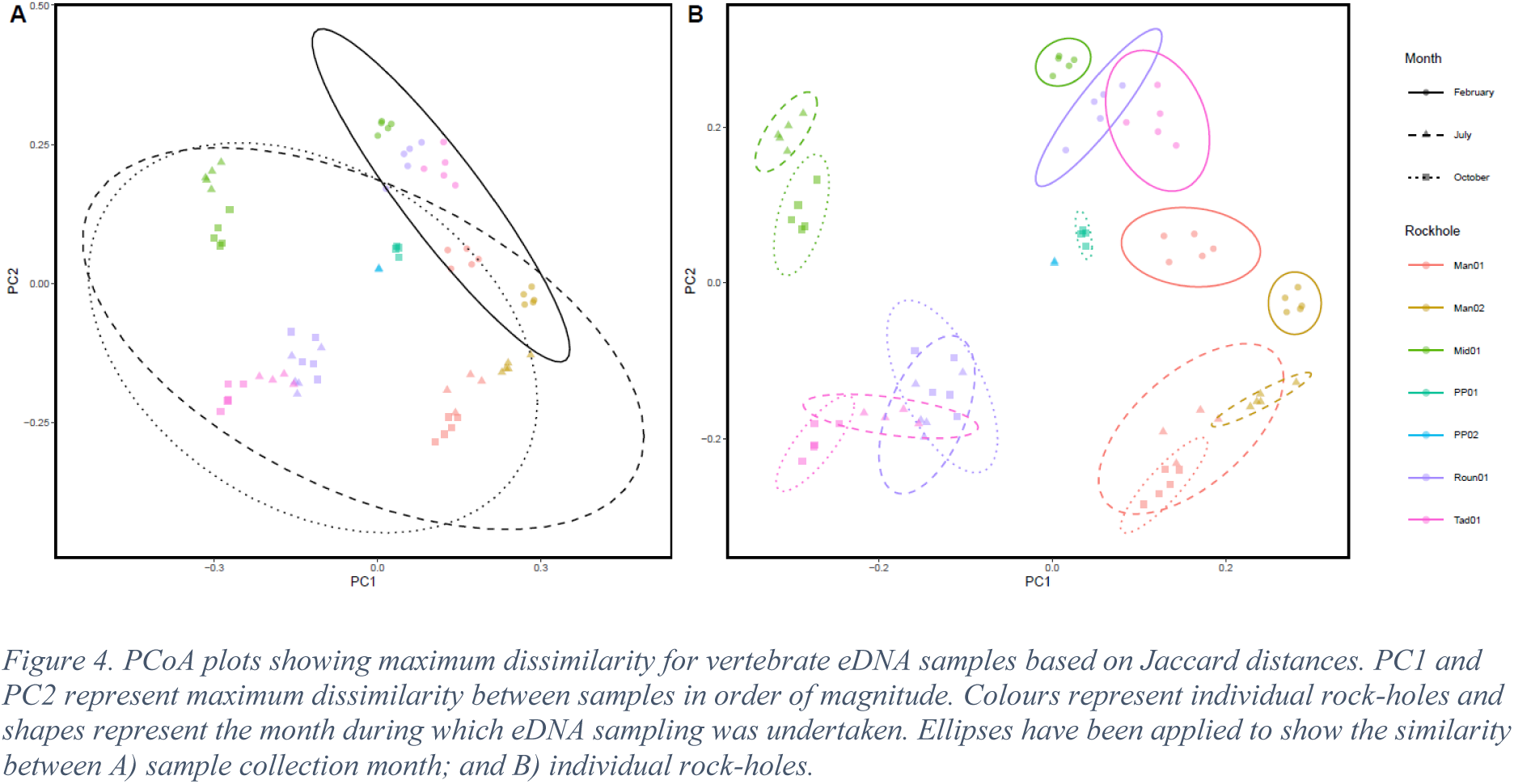
PCoA plots showing maximum dissimilarity for vertebrate eDNA samples based on Jaccard distances. PC1 and PC2 represent maximum dissimilarity between samples in order of magnitude. Colours represent individual rock-holes and shapes represent the month during which eDNA sampling was undertaken. Ellipses have been applied to show the similarity between A) sample collection month; and B) individual rock-holes.

## Discussion

Here we contribute to an emerging narrative regarding the detection of vertebrate visitors to arid-lands freshwater ecosystems using eDNA methods as compared to the traditional wildlife camera trap monitoring method (Farrell et al. 2022, Croose et al. 2023, Johnson et al. 2023 McDonald et al. 2023). Our results demonstrate for the first time the efficacy of eDNA as a complementary tool suitable for monitoring both native and invasive vertebrate visitation to arid-lands rock-holes. We observed that vertebrate eDNA was successfully recovered from freshwater samples collected from rock-holes and complementary to the use of wildlife cameras for monitoring mammals and amphibians, although for birds and reptiles a combined approach is likely required to capture the complete detectable community. We discuss our observations further below.

Anecdotally, rock-holes are known to be used by native vertebrates in Australian arid regions, but reliable scientific data is rare (McDonald et al. 2023). Our findings provide new evidence that the freshwater rock-holes in the Gawler bioregion are of conservation value due to their use by local native vertebrates. We observed a suite of 21 vertebrate species using eDNA water samples from seven rock-holes (Figure 2), capturing 11% of vertebrates known from this region (Commonwealth of Australia 2015a). The community composition of vertebrates recovered using eDNA was similar to that recovered through wildlife camera trapping at the same locations (Figure 2). The taxa recorded visiting at the highest frequency (number of individual photographs recorded) using wildlife camera traps (i.e., macropods and emus), were also amongst those detected in most eDNA samples (70-85%). The rarer species such as the southern hairy-nosed wombat (*Lasiorhinus latifrons*) and feral cats (*Felis catus*) were detected in the fewest eDNA samples (15%).

Invasive species are known to inhabit the Australian arid-lands (Commonwealth of Australia 2015a). Damage caused by such species includes predation inconsistent with pre-European levels, competition for resources including water, impacts to vegetation composition and structure, toxicity and disease transmission (Brim Box et al. 2019, Feit et al. 2020, Jansen et al. 2022, Stobo-Wilson et al. 2022). Here we observed that invasive species known to cause ecological harm in the region (Commonwealth of Australia 2008a, Commonwealth of Australia 2008b, Commonwealth of Australia 2015b, Jansen et al. 2022) accounted for 20% of the species detected with eDNA. We observed feral goats to be detected in the greatest number of samples in addition to cats, foxes, and rabbits (Figure 2). Significantly, data from eDNA revealed complimentary results to camera trap data except for three invasive mammal species detected using eDNA which were not detected using traditional wildlife cameras (i.e., cows, sheep, and dogs). While we approach these results with caution since these species are known to have a higher likelihood of occurring through sequence contamination than other taxa present (Zhang et al. 2023), the results hold promise for improved results on the traditional camera trap approach. Given that early detection of invasive species using eDNA metabarcoding is an emerging discipline that shows promise (Rishan et al. 2023, Salis et al. 2023, Zhang et al. 2023) and the findings of our study indicate a strong proof of concept that these methods are extendable into semi-arid environments for survey of local vertebrate fauna.

Our results demonstrated a high degree of similarity between replicates at individual rock-holes indicating consistency and validation between replicates. In contrast, eDNA showed highly distinct communities between separate rock-holes (Figure 4). Whilst this suggest that each rock-hole is used by a distinct suite of taxa, it is likely that an increased sample size would shed light on this aspect of rock-hole use. Furthermore, eDNA was variable in its detection consistency when compared to vertebrate visitation recorded from wildlife cameras throughout the same period. We observed that 80% of mammals detected using wildlife cameras were also recovered using eDNA, but only 11% of birds and 25% of reptiles were similarly recovered. Other studies report similar results regarding taxon-specific detection, with Zhang et al. (2023) demonstrating that water birds are easier to detect using eDNA than forest birds. Reasons for inconsistent or reduced detection of birds and reptiles are four-fold. First, eDNA presence/absence may be linked to taxon-specific physiology or behaviour where less frequent visitation at rock-holes by birds and reptiles may be driving decreased detectability (Hedges unpublished data). Second, reduced detectability may also be due to lower rates of eDNA deposition by birds. It is known that keratinised beaks and feathers are poor sources of eDNA (Turcu et al. 2023) and contact with water with these tissue types is likely to impact delivery of DNA to the substrate. Third, assay choice may also impact detection of species. Whilst we observed successes in recovery of mammal sequences, and comparison with camera trapping data suggest that the *16S* vertebrate assay used here is suitable for detecting a broad range of species in the arid-lands. The technique was notably less successful at detecting birds and reptiles than wildlife camera trapping. This decrease in detection success may suggest the *16S* vertebrate assay is less well suited for detecting birds and reptiles than for mammals. Finally, public DNA reference libraries typically provide both high quality and quantity of sequences for mammal fauna, but are less comprehensive for reptile and bird fauna (see further below).

Vertebrate visitation to rock-holes at HNR varied spatially based on our eDNA data (Figure 4). We observed rock-holes within a granite outcrop to be more similar to one another than to those on other outcrops, with absences of multiple macropod species and emus responsible this variability (Figure 4). It is possible that elevation impacted species access to rock-holes as the two outcrops here vary by approximately 90 metres in elevation, with significantly steeper slopes also being present at the Pretty Point outcrop than at the Photopoint outcrop (Table 1). However, access limitations at multiple outcrops and an absence of water in some rock-holes during sampling times meant replication was insufficient to accurately test the impact of outcrop location on vertebrate visitation. Our findings were consistent with Hedges (2023), where it was shown that various rock-holes attracted unique vertebrate communities and that rock-hole elevation was likely responsible for this dissimilarity, as at higher elevations (>250 m) they are less accessible to taxa which are unable to climb the steep slopes characteristic of the Gawler bioregion. McDonald et al. (2023) also explored visitation to similar rock-holes and suggested that some taxa may avoid certain rock-holes due to increased risk of injury or predation. Similarly, Votto et al. (2022) recorded visitation to arid-lands waterholes, and found that visitation rates were impacted by fringing vegetation.

Based on our eDNA results, vertebrate visitation to freshwater rock-holes at HNR also varied temporally over the year (Figure 4). Whilst communities recovered at the beginning (July) and end (October) of the winter wet period were relatively similar, both communities were distinct from those recovered during the summer wet period in February (Figure 4). Native species’ (primarily macropods, emus, crows, and ravens) dependence on freshwater available at rock-holes was observed to increase with time elapsed since rainfall events, with rock-holes increasing in their resource value with every successive week without rain, although this relationship was not observed for invasive species (primarily feral goats) (Hedges 2023). Freshwater visitation by vertebrates is often temporally variable (Eliades et al. 2022, Kassara et al. 2023) and, as such, it is likely that future biomonitoring efforts using eDNA will need to take time of sampling into consideration, as periods when visitation peaks are likely to provide the greatest indication of resource use (Sales et al. 2020b).

Overall, we show that the efficacy of eDNA as a tool for monitoring vertebrate visitation at freshwater systems and provide a foundation for further use of the technique in monitoring semi-arid and arid freshwater ecosystems. We consider visitation frequency to be a likely a major factor in determining eDNA signal (Ushio et al. 2017, Sales et al. 2020b) and future research would benefit from testing the relationship between visitation frequency and eDNA concentration. Given that the detection of the complete suite of vertebrate fauna at arid-lands rock holes is variously detected with both camera trap and eDNA methods, we suggest a complimentary approach using a combination of traditional and eDNA techniques may be favourable when characterising vertebrate communities accessing desert freshwater habitats. For birds and reptiles, we emphasise the use of more targeted freshwater eDNA approaches such as species-specific assays (e.g., White et al. 2020).

### Implications for management at Hiltaba Nature Reserve

The frequent occurrence of invasive species (primarily feral goats) in our eDNA metabarcoding results suggest these species are capitalising on an unmanaged freshwater resource and that rock-hole management has the potential to influence invasive species activity throughout the Gawler bioregion. Managing their access to this resource may allow for population suppression and improve native species outcomes for the broader landscape. Invasive species visitation is also likely to negatively impact the rock-holes themselves, the communities of native species reliant on these water sources, and the quality of the water therein. Ungulates can cause degradation of Australian freshwater ecosystems through disturbance and the input of faeces, which dramatically increases nutrient loads (Doupe 2010, Brim Box et al. 2016). Efforts to limit invasive species access to the rock-holes are therefore likely to benefit both invasive species management programs over a wider area, alleviate their impacts in the Gawler bioregion, and benefit the rock-hole ecosystem itself. Future and broader rock-hole eDNA studies may provide further clarity on the rock-holes that are most frequently attended by invasive species, facilitating more effective culling in the region.

### Environmental DNA as a biomonitoring tool

Our results demonstrate the efficacy of eDNA as a tool for monitoring freshwater rock-holes in Australia’s semi-arid and arid lands, performing well compared to more traditional survey methods such as typical human-led field surveys and wildlife camera trapping. Multiple advantages exist in the use of eDNA as a monitoring tool. The collection of freshwater eDNA samples requires no specialised scientific training and can therefore be carried out by landholders, conservation workers, volunteers, and other stakeholders with ease (Prié et al. 2021, 2023). Whilst specialised equipment is required for on-site filtering of samples, it is relatively affordable and very quickly becoming unnecessary (Bessey et al. 2021, van der Heyde et al. 2023). Passive freshwater eDNA collection has been trialled in similar rock-holes elsewhere in Australia through the submergence of filter papers (McDonald et al. 2023), and removes the filtration step used here, further cutting costs and effort related to sample collection. However, communities recovered from passively collected samples have been shown to capture only a subset of freshwater communities compared to those recovered from filtered samples (McDonald et al. 2023). Once filtered, samples may be easily stored in a conventional freezer, and can be transported on ice; alternatively, a range of ambient preservation techniques are being explored to remove the need for cold-chain transport. Sequencing costs can be high, particularly for eDNA samples which are sensitive to contamination (Zhang et al. 2023). However, it is likely that costs will continue to decrease as advancements are made in the field and eDNA becomes a more widely applied method (Rishan et al. 2023).

Limitations in the availability of high-quality, taxonomically-verified reference libraries are an obstacle that will need to be overcome before eDNA can be widely applied as a biomonitoring tool, including at the ephemeral rock-holes of the Gawler Bioregion (Beasley-Hall et al. 2023). Remote semi-arid and arid Australia is broadly understudied and for many taxonomic groups, including some vertebrates, very little work has been done to generate sequences corresponding to barcoding regions commonly used in eDNA studies. Greater sampling and sequencing of semi-arid and arid Australian taxa is therefore a critical research priority and would facilitate more effective use of eDNA methods for monitoring the biodiversity of those regions. Primers specifically designed for detection and identification of terrestrial mammals will improve the efficacy of such studies (Ushio et al. 2017, Leempoel et al. 2020). Assays targeting birds and reptiles may allow for greater success in detection of these taxa from freshwater samples (Neice and McRae 2021, Mousavi-Derazmahalleh et al. 2023). Furthermore, an improved understanding of how organism size, physiology, and ecology impacts eDNA deposition would allow greater confidence in determining true species absences in instances when eDNA has not been recovered for a particular taxon.

## Conclusions

Our findings demonstrate the viability of freshwater eDNA metabarcoding as a method for monitoring vertebrate visitation to granite rock-holes in the Gawler bioregion, with both native and invasive species successfully detected. We found that vertebrate visitation is variable among rock-holes at HNR, with different outcrops displaying different communities. Furthermore, vertebrate visitation was variable throughout time, with communities recovered during different months being distinct from one another. With eDNA metabarcoding, we recovered sequences that indicate that invasive species are visiting rock-holes throughout the region and are likely impacting them and their value as a freshwater resource for vertebrates of conservation interest. We recommend that future studies improve reference databases to allow better taxonomic assignment within eDNA datasets.

